# Detection of Pancreatic Cancer Using a Methylation-Specific PCR-Based Multi-Cancer Early Detection Test

**DOI:** 10.64898/2026.05.27.728292

**Authors:** Hanh T. Pham, Kimberly J. Bussey, Marc M. Oshiro, Matthew Rounseville, Michael Moses, Alejandro Zulbaran-Rojas, Vincent Nguyen, Richard A. Bernert, Joshua Routh, George Watts, Geoffrey D. Block, William E. Fisher, Mark A. Nelson

## Abstract

**Context:** Pancreatic ductal adenocarcinoma (PDAC) is an aggressive malignancy often diagnosed at advanced stages due to the lack of early clinical symptoms. DNA methylation alterations arise early in PDAC tumorigenesis and may serve as promising biomarkers for blood-based cancer detection.

**Objective:** To evaluate the performance of EPISEEK, a laboratory-developed blood-based multi-cancer early detection (MCED) assay, for detecting PDAC across disease stages.

**Design:** A retrospective cohort study included 97 patients with stage I–IV PDAC and 201 asymptomatic healthy controls. Sensitivity, specificity, area under the curve (AUC), and stage-specific performance were assessed. EPISEEK-MCED performance was also compared with CA 19-9 alone and in combination with CA 19-9.

**Results:** EPISEEK-MCED classified 65 of 97 PDAC cases as positive, corresponding to an observed sensitivity of 70.1% (95% CI, 60.3% - 78.3%) at 99.5% specificity. The assay demonstrated strong discrimination between PDAC cases and healthy controls, with an AUC of 0.916 (95% CI, 0.88 - 0.952). Sensitivity increased with advancing stage while remaining substantial in early-stage disease, measuring 53.6% for stage I and 65.1% for stage II PDAC, 100% for stage III and 94.7% for stage IV. Across stages, EPISEEK-MCED outperformed CA 19-9 alone, particularly in early-stage disease. Combined analysis of EPISEEK-MCED and CA 19-9 further improved detection performance, achieving sensitivity of 57.1% and 81.4% for stage I and II, respectively.

**Conclusions:** EPISEEK-MCED demonstrated high specificity and sensitivity for PDAC detection across disease stages, including early-stage disease. Combining EPISEEK-MCED with CA19-9 further improved performance, supporting its clinical utility for PDAC detection.

## INTRODUCTION

Pancreatic cancer is projected to be the second leading cause of cancer deaths in the United States by 2040 ^1,2^. Hence, reliable early detection approaches for pancreatic cancer are needed. The importance of early detection is underscored by the fact that most pancreatic cancers are diagnosed in late stages (approximately 80% per SEER data or as high as 85% in unscreened populations in the CAPS5 surveillance study), with a 5-year survival rate as low as 3% for individuals with metastatic disease ^1,3^. Whereas the 5-year survival rate for localized disease is 44% ^2^.

A promising approach for early detection and monitoring of pancreatic cancer is liquid biopsy, which analyzes cancer-related material, such as DNA, from blood samples ^4,5^. Some methods focus on detecting tumor-specific biomarkers in the cell-free DNA (cfDNA), particularly circulating tumor DNA (ctDNA), which varies by tumor type and disease stage. Methylated DNA markers are often more informative than mutations and can be specific to tumor origin ^6,7^. Since tumors typically exhibit many aberrantly methylated regions, analyzing multiple genomic loci using methods like methylation-specific qPCR can enhance detection sensitivity. Therefore, identifying tumor-specific DNA methylation in cfDNA offers potential for early pancreatic cancer.

A liquid biopsy test (termed EPISEEK Multicancer Early Detection Test, EPISEEK-MCED) has been developed and validated as a laboratory developed test performed at Precision Epigenomics Inc in Tucson, Arizona assessing aberrant DNA methylation ^8^. Previous studies evaluated the ability of a cfDNA methylation biomarker panel used in a multi-cancer early detection assay to distinguish metastatic pancreatic cancer cases from benign pancreatic cyst patients without dysplasia on histology, as well as to monitor tumor dynamics following chemotherapy ^9^. The biomarkers were able to distinguish malignant cases from benign disease with high sensitivity and specificity (AUC = 0.999) ^9^. In addition, the biomarkers detected a consistent decline in cancer derived cfDNA in serial samples from patients undergoing chemotherapy ^9^. This suggests EPISEEK-MCED liquid biopsy assay may aid in managing pancreatic cancer. Given the fact that there is a lack of early detection approaches available for pancreatic cancer, the goal of this study was to assess the ability of the EPISEEK-MCED test to detect early-stage pancreatic cancer. We found that the EPISEEK-MCED test could detect all stages of pancreatic cancer. These findings demonstrate the potential utility of the EPISEEK-MCED test for the detection of pancreatic cancer across all stages, including early-stage disease, underscoring its promise as a non-invasive diagnostic tool to address the current gap in early detection strategies.

## METHODS

### Participants

This was a retrospective study. For the cancer cohorts, blood samples from Baylor College of Medicine (BCM) were collected in Vacutainer EDTA from 64 patients diagnosed with Stage I, II, III, or IV PDAC, as well as from 21 patients with various non-PDAC pancreatic lesions that included both benign and pre-malignant pathologies.

These patients underwent pancreatic resection, and blood was collected pre-operatively. A few specimens had been banked for as long as 20 years, but the majority (51/64) were collected after 2020, with available plasma volumes ranging from 0.4 to 2.0 mL (mean volume 1.43 mL). In addition, Streck tube blood specimens were collected from 29 pancreatic cancer patients from the University of Arizona, with 0.9 to 2 mL (mean volume 1.05 mL) of banked plasma for examination. An additional 4 samples collected in Streck tubes were purchased from Reference Medicine. Institutional Review Board approval from both institutions was obtained prior to the study initiation and all participants provided written informed consent.

The control reference cohort consisted of 201 asymptomatic individuals with no known history of malignancy with demographic information available in Table 1. Blood from these participants was collected in Streck cfDNA BCT tubes to preserve cell-free DNA integrity. Data from this control group were used to establish the baseline reference range for each methylation biomarker included in the assay and to develop and train the multivariate qualitative classifier used for cancer detection.

**Table 1.**
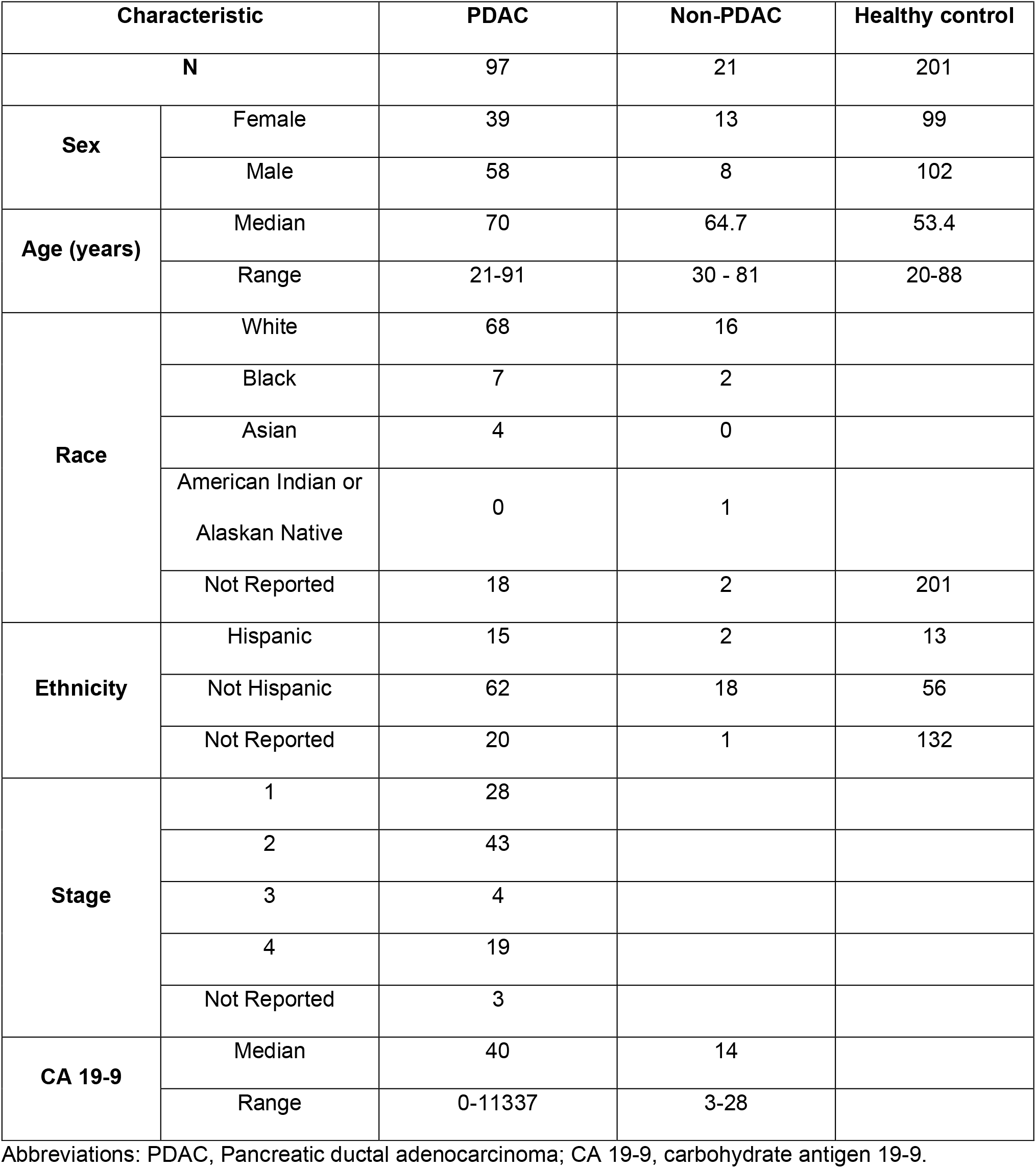
Baseline Demographic and Clinical Characteristics of PDAC and Non-PDAC Study Cohorts.

### Assay workflow

The EPISEEK-MCED assay was conducted under unblinded and blinded conditions in the BCM cohort. The Arizona cohort was conducted in blinded conditions. The clinical laboratory workflow for the EPISEEK-MCED (University of Arizona) assay follows methods previously reported ^10, 11^. After blood collection in Streck cfDNA BCT tubes (La Vista, NE) or Vacutainer EDTA tubes, plasma was processed as previously described^10^. In brief, plasma was separated by two-step centrifugation and stored at -80 °C when needed. cfDNA was isolated from 2 mL plasma using the MagMax cfDNA Isolation Kit (Applied Biosystems). The cfDNA underwent bisulfite conversion using the EZ-96 DNA Methylation-Gold MagPrep (Zymo Research), then was subjected to 2-step polymerase chain reaction (PCR). A two-step quantitative PCR (qPCR) protocol consisting of 15 cycles of multiplex pre-amplification, followed by 50 cycles of locus-specific qPCR, as previously established using primers targeting 10 cancer-associated regions selected for their ability to detect a broad range of common carcinomas^11^. The design of corresponding qPCR amplicons was previously reported, ^9,10,12^ and qPCR was performed on QuantStudio 7 Pro Dx real-time PCR systems (Thermo Fisher Scientific). Cycle threshold (Ct) values were generated using QuantStudio software, and methylation scores were calculated by integrating Ct values from target loci and internal controls. Results were generated using an interpretive algorithm developed for the EPISEEK-MCED V. 1.02 assay.

### Data Analysis

DNA methylation-based disease classifiers have emerged as powerful tools for early cancer detection ^13,14^. An interpretive algorithm, the EPISEEK-MCED V.1.02, was developed using Ct values from blood samples from 201 asymptomatic individuals with no known history of cancer. These data were used to establish a reference range for each biomarker and to train the multivariate qualitative EPISEEK-MCED V1.02 classifier. The EPISEEK-MCED V1.02 classifier was then applied to the qPCR data generated from the analysis of the pancreatic biospecimens described above.

For each sample, the methylation score represents the negative log-odds of the second-highest methylation probe value belonging to the distribution of probe values observed in 201 control plasma samples. Ct values were normalized to the average value of the internal control probes. These values were then fitted to the probe-specific distribution, and the negative log-odds were computed. The probes were ranked by score, and the second-highest score was compared to a test-specific cut-off value that yields a 99.5% specificity to be called positive or negative classification.

### Statistics

All statistical analyses were conducted in R (version 4.5.1). Sensitivity, specificity, and receiver operator curve (ROC) analyses were performed using the pROC (version 1.19.0.1), rstatix (version 0.7.3), and fastR2 (version 1.2.5) packages. Cox proportional hazard analysis adjusting for stage, age, race, and ethnicity and log-rank evaluation of Kaplan-Mier curves was done using the survival (version 3.8-3) and survminer (version 0.5.1) packages. ANOVA analysis followed by Tukey Post-Hoc was used to determine the differences between control, non-PDAC diagnoses, stage I/II PDAC, and stage III/IV PDAC.

## RESULTS

To evaluate the ability of EPISEEK-MCED to detect early-stage PDAC, we retrospectively analyzed the plasma of 97 participants with PDAC and 21 with non-PDAC benign or pre-malignant pancreatic lesions. Plasma was collected in either Streck or purple top EDTA tubes. The demographics of the patients are given in Table 1.

Receiver operating characteristic (ROC) analysis confirmed the strong discriminatory performance of the EPISEEK-MCED assay for differentiating PDAC cases from controls (Figure 1). The assay achieved an area under the curve (AUC) of 0.915 (95% CI, 0.88 - 0.952), indicating robust separation between PDAC and control samples across the range of classification thresholds. At the predefined operating point corresponding to 99.5% specificity, the observed assay sensitivity across all PDAC stages was 70.1% (95% CI, 60.1% - 78.3%).

**Figure 1.**
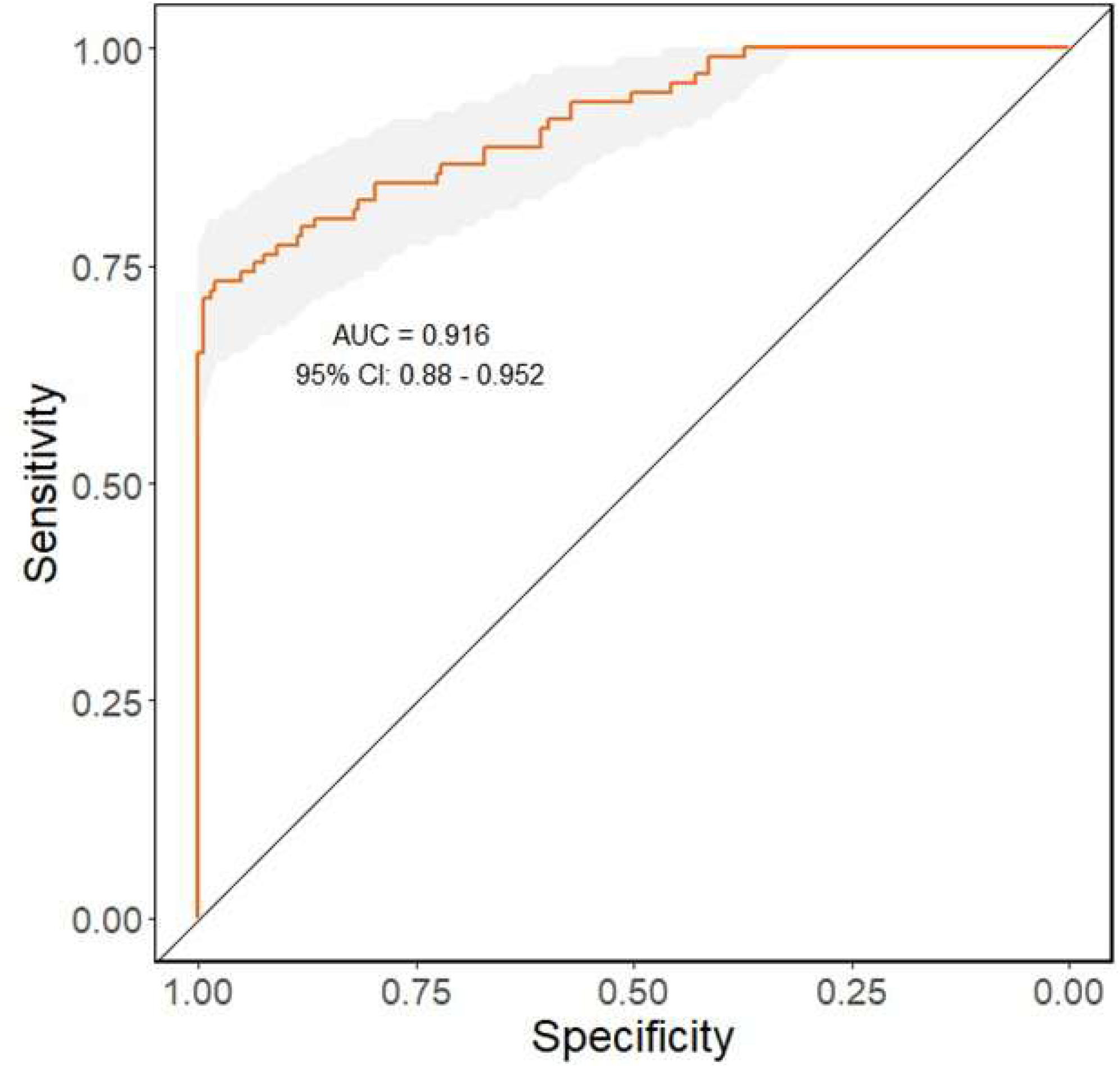
ROC Curve for EPISEEK-MCED classifier. ROC analysis on the 84 pancreatic cancer cases and 201 controls demonstrates an AUC of 0.916 (95% CI 0.88 – 0.952), and an accuracy of 89.9% (95% CI, 85.9–92.9%). Abbreviations: ROC, Receiver-Operator Curve; MCED, Multicancer Early Detection; AUC, area under the curve.

The distributions of EPISEEK-MCED methylation scores across healthy controls and PDAC cases, stratified by disease stage, are shown in Figure 2. ANOVA followed by pairwise comparisons revealed statistically significant differences between healthy controls, early-stage PDAC, and late-stage PDAC (Tukey post-hoc, all comparisons *P* < .001). Late-stage PDAC (stage III/IV) exhibited higher methylation scores with greater separation from both control and early-stage groups. The EPISEEK-MCED test was able to detect all stages of pancreatic cancer.

**Figure 2.**
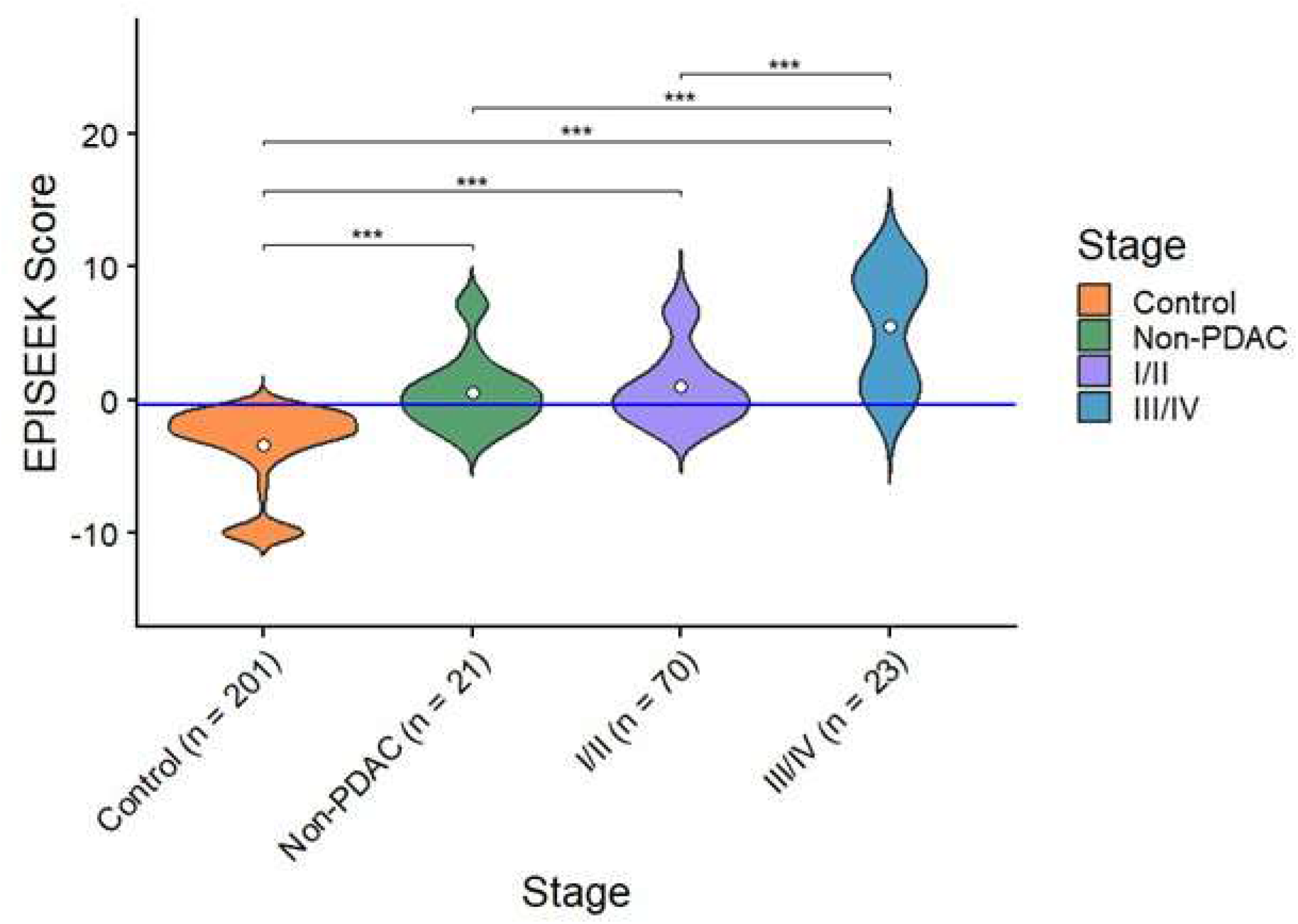
Distribution of EPISEEK-MCED Methylation Scores Across Control, Non-PDAC lesions, and PDAC Disease early stage I/II and late stage III/IV. Violin plots illustrate the distribution of EPISEEK-MCED methylation scores among healthy controls (n = 201), non-PDAC pancreatic lesions (n = 21), early-stage PDAC (stage I/II; n = 70), and late-stage PDAC (stage III/IV; n = 23). The width of each violin reflects the density of observations, with white dots indicating median values. The horizontal blue line represents the predefined assay threshold. Data were analyzed by ANOVA followed by a Tukey Post-hoc test which controls for multiple comparisons. * *P* < .05, ** *P* < .01, *** *P* < .001. Overall, EPISEEK-MCED methylation scores increased with advancing disease stage, demonstrating clear separation among control, non-PDAC, early-stage PDAC, and late-stage PDAC groups. No statistically significant difference observed between non-PDAC lesions vs early-stage diseases. Abbreviations: PDAC, Pancreatic ductal adenocarcinoma.

In addition to the PDAC cohorts, a set of at-risk pancreatic lesions were tested. These included 8 subjects with intraductal papillary mucinous neoplasms (IPMN), 4 with cystadenomas, 1 bile duct stricture, 1 ampullary metaplasia, 1 chronic pancreatitis, 2 pseudopapillary neoplasms, 2 pancreatic retention cysts, 1 intrapancreatic ectopic spleen, and 1 pseudocyst. We found 4 out 8 (50%) of the IPMN positive and 3 out 4 of the cystadenomas positive. In addition, the ampullary metaplasia, pseudopapillary neoplasms, and a benign retention cyst were positive. Other pancreatic lesions were negative (Table 2).

**Table 2.** Sample size and Detection Rates of the EPISEEK-MCED Assay in Non-PDAC at risk Pancreatic Lesions.

| Lesion type | Number tested | Number positive | Positivity (%) |
| --- | --- | --- | --- |
| Intraductal papillary mucinous neoplasms (IPMN) | 8 | 4 | 50% |
| Cystadenoma | 4 | 3 | 75% |
| Pseudopapillary neoplasm | 2 | 2 | 100% |
| Ampullary metaplasia | 1 | 1 | 100% |
| Pancreatic retention cyst | 2 | 1 | 50% |
| Bile duct stricture | 1 | 0 | 0% |
| Intrapancreatic Ectopic Spleen | 1 | 0 | 0% |
| Chronic Pancreatitis | 1 | 0 | 0% |
| Pseudocyst | 1 | 0 | 0% |

Abbreviations: MCED, Multicancer Early Detection; PDAC: Pancreatic ductal adenocarcinoma IPMN, Intraductal papillary mucinous neoplasms.

For those samples for which we had CA 19-9 values, we compared the performance of EPISEEK-MCED to CA 19-9, calling samples with CA 19-9 values greater than 37 positives (Table 3). EPISEEK-MCED had improved performance, particularly in early-stage PDAC (Table 3). When we combined EPISEEK-MCED with CA 19-9, we had improved sensitivity, particularly in stage I cancers, where sensitivity rose from 35.3% with CA 19-9 alone to 53.6% with EPISEEK-MCED alone and 57.1% combined.

**Table 3.** Stage-Specific Sensitivity of EPISEEK-MCED, CA-19-9, and the Combined Assay in PDAC detection.

| **Stage** | **EPISEEK-MCED Sensitivity** | **EPISEEK-MCED 95% CI** | **CA 19-9 Sensitivity** | **CA 19-9 95% CI** | **EPISEEK-MCED + CA 19-9 Sensitivity** | **EPISEEK-MCED + CA 19-9 95% CI** |
| --- | --- | --- | --- | --- | --- | --- |
| All | 70.1% | 60.3% - 78.3% | 52.1% | 38.1% - 63.6% | 78.3% | 69.0% - 88.4% |
| 1 | 53.6% | 35.8% - 70.4% | 35.3% | 17.3% - 58.9% | 57.1% | 39.1% - 73.4% |
| 2 | 65.1% | 50.0% - 77.6% | 58.8% | 43.0% - 73.6% | 81.4% | 67.0% - 90.4% |
| 3 | 100 % | 45.0% - 100% | - | - | 100 % | 45.0% - 100% |
| 4 | 94.7 % | 73.2% - 100% | 75.0% | 29.0% - 96.0% | 94.7 % | 73.2% - 100% |

Abbreviations: MCED, Multicancer Early Detection; PDAC: Pancreatic ductal adenocarcinoma; CA 19-9, carbohydrate antigen 19-9.

We explored the relationship between score, demographic factors, and survival. A Cox proportional hazard analysis failed to demonstrate a significant relationship between score and survival after controlling for age, sex, race, and ethnicity (data not shown).

Linear regression analysis and logistic regression analyses of the relationship between score and demographic factors yielded interesting results. In the linear regression analysis, which uses the numerical value of the score as the dependent variable, being male is associated with a 1.5-unit lower score (*P* = .0476). In contrast, being Hispanic or Latino is associated with a 3.15-unit increase in score (*P* = .001). However, this relationship does not translate into a change in the odds ratio when we dichotomize the dependent variable to positive and negative calls (data not shown).

## DISCUSSION

Pancreatic cancer remains a highly lethal malignancy largely due to the absence of early clinical symptoms and the consequent diagnosis at advanced stages when therapeutic options are limited. Epigenetic alterations, particularly aberrant DNA methylation, arise early in pancreatic tumorigenesis and can be detected in circulating cell-free DNA, offering a potential avenue for noninvasive early detection. Despite this biological rationale, there is currently no routinely recommended screening strategy for pancreatic cancer. In this context, EPISEEK, a clinically validated MCED assay that interrogates ten differentially methylated genomic regions altered across more than 65 cancer types, represents a promising approach. The present study evaluated the performance of the EPISEEK-MCED V1.02 assay for the detection of pancreatic ductal adenocarcinoma, providing insight into its potential utility for pancreatic cancer detection. The assay achieved an overall AUC of 0.916, demonstrating high discriminatory power against asymptomatic controls, with a sensitivity of 70.1% at 99.5% specificity across all disease stages. Notably, robust performance was maintained in early-stage disease, with sensitivity of 53.6% (95% CI: 35.8% - 70.4%) and 65.1% (95% CI: 50.0% - 77.6%) observed for stage I and stage II PDAC, respectively; sensitivities of 100% (95%CI: 45% - 100%) in stage III and 94.7% (95% CI: 73.2% - 100%) stage IV cases were seen.

Compared with existing assays developed for PDAC, mutation-based ctDNA assays, particularly those targeting KRAS mutation, have shown limited sensitivity in early-stage disease ^15–17^. Although *KRAS* mutations are present in over 90% of PDAC cases, assays detecting mutant *KRAS* in ctDNA obtained sensitivity below 40% in early-stage patients ^18,19^. This reflects two biological limitations in the early stage of PDAC: the inherently restricted ctDNA shedding of early-stage pancreatic tumors and the substantial hepatic filtration of tumor-derived DNA before it reaches the peripheral circulation. Multi-analyte blood tests, such as CancerSEEK or CancerGuard, have shown the ability to detect early-stage PDAC, with a reported sensitivity of about 42.9% and 75.0% for stage II at 99.5% specificity ^20,21^. Methylation-based sequencing approaches such as Galleri multi-cancer early detection NGS-based platform, achieved PDAC sensitivities around 61.9% for stage I and 60% for Stage II PDAC at 99.5% specificity ^22–24^. Pancreatic cancer detection using 5-hydroxymethylcytosine signals has been reported to have a sensitivity of 68.3% and a specificity of 96.9% for stage I/II PDAC ^25,26^. In this context, the performance of EPISEEK-MCED compares favorably, especially given its PCR-based workflow and lower input than sequencing requirements ^27^. These results place EPISEEK-MCED among the most effective blood-based assays for high discrimination against asymptomatic controls with 95% confidence intervals that overlap those reported for tests like Galleri and CancerGuard ^21–23^.

Serum carbohydrate antigen, CA 19-9, remains the only tumor marker primarily used to monitor disease burden and treatment response ^28,29^. However, CA 19-9 demonstrates limited sensitivity in early-stage disease, typically 30-50%. CA 19-9 specificity is also reduced because its level can rise in non-malignant or benign conditions, producing false positive results. Additionally, CA 19-9 is not expressed in Lewis antigen-negative individuals, which reduces sensitivity for PDAC detection in that subgroup ^28,30^. In our cohort, CA 19-9 detected only 31.3% of stage I PDAC cases, whereas EPISEEK-MCED detected 55.6%, suggesting EPISEEK-MCED identified a substantial proportion of CA 19-9-negative cases, highlighting its potential to address a critical sensitivity gap in pancreatic cancer detection. In addition, among CA 19-9-positive cases, the EPISEEK-MCED methylation score provided tumor-specific confirmation, which may help to reduce diagnostic ambiguity arising from non-malignant elevations of CA 19-9.

The combined use of EPISEEK-MCED and CA 19-9 further improved detection sensitivity, reaching 57.1% for stage I and 81.4% for stage II disease, exceeding the performance of CA 19-9 alone and comparable to that reported for multi-cancer sequencing-based platforms. Our findings are consistent with prior studies demonstrating improved sensitivity when CA 19-9 is combined with methylation- or ctDNA-based biomarkers ^31,32^. Collectively, these results suggest that a combined methylation and CA 19-9 strategy offers complementary strengths that may enhance early detection and diagnostic confidence in PDAC and other upper gastrointestinal malignancies.

Demographic analyses identified modest associations between methylation score and sex or Hispanic/Latino ethnicity; however, these effects did not influence the final test call, indicating robustness of the EPISEEK-MCED current thresholding approach. Similar findings have been reported across multiple cfDNA methylation-based cancer detection platforms, including Galleri, PanSeer, DELFI-methylation hybrid models, and TruSight Oncology ctDNA methylation classifiers ^29,33,34^, suggesting that while baseline methylation patterns may vary by demographic group, such variation does not meaningfully affect assay performance or clinical classification.

Previous reports have demonstrated the diagnostic efficacy of this assay in distinguishing PDAC from benign pancreatic cyst cohorts, with high sensitivity and specificity observed under well-defined conditions ^9^. In the current study, from 21 disease controls, 11 patients yielded positive results with the EPISEEK-MCED assay. It should be noted that the disease controls included in the current study largely comprise precancerous or high-risk pathological conditions such as IPMN, Cystadenoma and Pseudopapillary Neoplasm. While not all pancreatic lesions progress to malignancy, IPMNs, characterized by intraductal papillary proliferation of mucin-producing epithelial cells, are well-recognized precursor lesions to PDAC. An important observation from this study is that the EPISEEK-MCED assay detected abnormal methylation signals in approximately 50% of individuals with IPMN (Table 2). Of the 4 IPMN with EPISEEK-MCED positives, there was 1 high grade dysplasia, the rest were low grade dysplasia (Table 2). All the negative IPMNs were low grade dysplasia. DNA methylation alterations are well-established early events in carcinogenesis, frequently preceding histologically detectable disease; it is plausible that the positivity detected in these cohorts reflects a biologically meaningful signal indicative of early epigenetic reprogramming associated with malignant transformation. This also highlights the challenges in differentiating non-PDAC pancreatic lesions from early-stage PDAC, a limitation that has also been reported in other liquid biopsy-based methods ^35–37^. These lesions may share overlapping epigenetic features or generate low-level cfDNA signals that complicate discrimination from early invasive disease ^38,39^. Future studies using the EPISEEK-MCED test in well-characterized IPMN cohorts could enable us to train our algorithms to detect precursor IPMNs with potential to progress to malignant lesions to enable closer surveillance of patients at risk. In addition, methylation-based assays combined with other diagnostic tools may enhance specificity for differentiating invasive PDAC from non-invasive precursor lesions at early PDAC stages.

Age-associated increases in DNA methylation are well-documented and statistically detectable and should be acknowledged as a general consideration in the application of methylation-based diagnostic biomarkers. In a previous study, healthy individuals spanning ages 25 to 75 years exhibited an approximately 2.5-fold increase in background methylation signal per marker across this five-decade range ^10^. Meanwhile, cancer patients demonstrated an overall 29-fold increase in marker methylation relative to healthy controls ^10^, representing a signal magnitude approximately 11.6-fold greater than that attributable to aging across the lifespan. Similarly, in this current study, a >20-fold increase in methylation was observed in both PDAC and non-PDAC groups compared to healthy controls. This pronounced differential indicates that marker positivity is primarily driven by pathological rather than epigenetic drift. Furthermore, the potential confounding effect of age had been addressed with an age-stratified analysis in the non-small cell lung cancer (NSCLC) cohort by restricting evaluation to the oldest tertile of controls (n = 16; 55-85 years), which was age-matched to the cancer cases ^10^. Within this subset, the EPISEEK-MCED panel maintained strong discrimination (AUC, 0.938), supporting robustness to age effects. Collectively, our findings demonstrate that although background methylation levels show a modest and predictable increase with advancing age, this effect does not meaningfully compromise the ability of the biomarker panel to distinguish cancer cases from healthy individuals, even within the oldest and most age-confounded stratum of the control population.

Despite these promising findings, several limitations in this study should be acknowledged. The absence of a matched control group limits direct assessment of clinical specificity in a real-world population. In addition, a subset of the BCM cohort was analyzed in an unblinded fashion. The small sample size, particularly for stage III and stage IV disease, reduces the precision of sensitivity estimates and underscores the need for additional late-stage PDAC samples. Also, limited cohort diversity further restricts the assessment of assay performance across broader populations. Future studies should include prospective evaluation in asymptomatic and high-risk populations, and validation in larger, more diverse cohorts to confirm and extend these findings.

## CONCLUSIONS

We successfully performed a retrospective study on a cohort of plasma samples collected from pancreatic cancer patients. EPISEEK-MCED demonstrates strong performance compared with existing liquid biopsy assays for PDAC, particularly improved sensitivity in early-stage disease and superior detection relative to CA 19-9 alone. Importantly, combining EPISEEK-MCED with CA19-9 significantly enhanced detection sensitivity. Collectively, these findings support the combined approach in earlier identification of resectable PDAC and improved clinical triage. Future studies should include prospective evaluation in asymptomatic and high-risk populations, and validation in larger, more diverse cohorts to confirm and extend these findings.

